# The accessory domains of Phospholipase D regulate its localization and activity in *Drosophila* photoreceptors

**DOI:** 10.1101/2024.09.22.614390

**Authors:** Amruta Naik, Padinjat Raghu

## Abstract

Phospholipase D (PLD) is activated during the response of animal cells to a large number of extracellular signals including growth factors, hormones and other extracellular stimuli. The product of PLD activity, phosphatidic acid (PA) is thought to act as a signalling molecule in cells. Since both the substrate, phosphatidylcholine and product phosphatidic acid (PA) are membrane anchored lipids, the precise localization and activation of PLD at specific locations in cells is critical. Here we report that that the precise localization of *Drosophila* PLD (dPLD) in photoreceptors is critical to maintain the structure of these cells during illumination. This localization of dPLD is dependent on its N-terminal PH domain; deletion of the PH domain (*dPLD*^Δ*PH*^) results in mis localization of the protein, loss of functional activity and *dPLD*^Δ*PH*^is unable to rescue the phenotypes of *dPLD* loss of function. By contrast, deletion of the PX domain (*dPLD*^Δ*PX*^) does not impact the localization of dPLD but enhances the functional activity of the protein. Thus the PH and PX domains of dPLD regulate its localization and activity respectively.

## Introduction

In addition to their function as structural components of cellular membranes, phospholipids also function as signalling molecules in eukaryotic cells. For example, the minor membrane lipid phosphatidylinositol is the substrate for the generation of its phosphorylated derivatives the phosphoinositides. Phosphoinositides can bind to proteins and regulate their activity and some such as PI(4,5)P_2_ can also be hydrolysed by phospholipase C (PLC) to generate second messengers that themselves can regulate cellular processes. In addition to PLC, the enzyme phospholipase D (PLD) has also been implicated in cell signalling. PLD that was first identified in plants, is a phospholipid-specific phosphodiesterase which hydrolyses phosphatidyl choline (PC) to generate membrane bound phosphatidic acid (PA) and free choline (Hanahan and Chaikoff, 1947). It has been known since the 1980s that PLD is a signal activated enzyme in mammalian cells. Many agonists including hormones and neurotransmitters activate PLD [reviewed in (Liscovitch et al., 2000)]. Interestingly many of these agonists also co-activate phospholipase C (PLC) which hydrolyses plasma membrane phosphatidylinositol 4,5 bisphosphate (PIP_2_) which is followed by increase in intracellular calcium [Ca^2+^]_i_ which relays downstream signalling. Studies in both yeast (Selvy et al., 2011) and *Drosophila* (Raghu et al., 2009a) have shown that PA formed by PLD activity exhibits properties of a signalling molecule that regulates downstream cellular processes. Genes encoding PLD are found in the genomes of a variety of organisms across the tree of life (Raghu et al., 2009b). While simpler eukaryotes code for a single gene encoding PLD activity, higher organisms for example mammalian genomes, contain 2 genes PLD1 and PLD2 that show PLD activity. The PLD protein has a conserved domain structure that includes the two HKD motifs which are important for catalytic activity (Frohman et al., 1999), the PH domain which is known to be important for membrane targeting (Sugars et al., 2002) and a polybasic PIP_2_ binding site that is required for enzymatic activity (Sciorra, 1999). Presumably, the localization and activity of PLD is regulated through the function of these additional domains acting in concert with the catalytic domains. However, the functional significance of these domains *in vivo* is not well studied.

A previous study (Thakur et al., 2016) has presented evidence that in *Drosophila* photoreceptors, where photons activate the G-protein coupled receptor rhodopsin leading to PLC activation, there exists a light induced dPLD activity. Loss of this dPLD activity does not directly affect the electrical response to light but during continued illumination results in progressive collapse of the apical microvillar membrane, a phenotype termed as a retinal degeneration (Thakur et al., 2016). During illumination, dPLD activity, regulates the recycling of endocytosed Rh1 from late endosomal compartment in an ARF1 and retromer complex dependent manner back to the plasma membrane (Thakur et al., 2016). In photoreceptors, dPLD is localized in the region of the sub-microvillar cisternae (LaLonde et al., 2005; Raghu et al., 2009a), a specialised sub-compartment of the smooth ER that forms a membrane contact site with the microvillar plasma membrane (Yadav 2016). However, the significance of the localization of dPLD to the SMC region in photoreceptors and the mechanisms regulating it remain unknown. In this study, we analysed the mechanisms by which dPLD is localized and the significance of this localization to the activity and function of the enzyme. We find, that while the PH domain is required for localising the enzyme at the SMC and this is localization is essential for the *in vivo* function of dPLD in photoreceptors. On the other hand, the PX domain appears to play no role in dPLD localization but appears to function as a negative regulator of dPLD activity. Thus, working together, the PH and PX domains of dPLD regulate it localization and activity to deliver physiological function *in vivo*.

## Results

### Overexpression of dPLD shows activity and light dependent retinal degeneration

We overexpressed of dPLD in the outer photoreceptors R1-R6 from ∼70% pupal development (*Rh1>dPLD*). This resulted in the progressive loss of rhabdomere structure as a function of age **[Fig. 1A]**. The retinal degeneration was dependent on illumination as *Rh1>dPLD* photoreceptors not exposed to illumination did not undergo retinal degeneration **[Fig. 1A]**. The degeneration phenotype was not observed upon overexpression of a lipase dead version of dPLD (*Rh1>dPLD ^K/R^*) su ggesting that the retinal degeneration was dependent on the catalytic activity of the enzyme [**Fig. 1A, B**]. The kinetics of degeneration increased with enhanced exposure to light (12hr L/D v CL) [**Fig. 1C, D**]. Together these findings show that under these conditions, the extent of retinal degeneration is directly proportional to dPLD activity.

**Figure 1:**
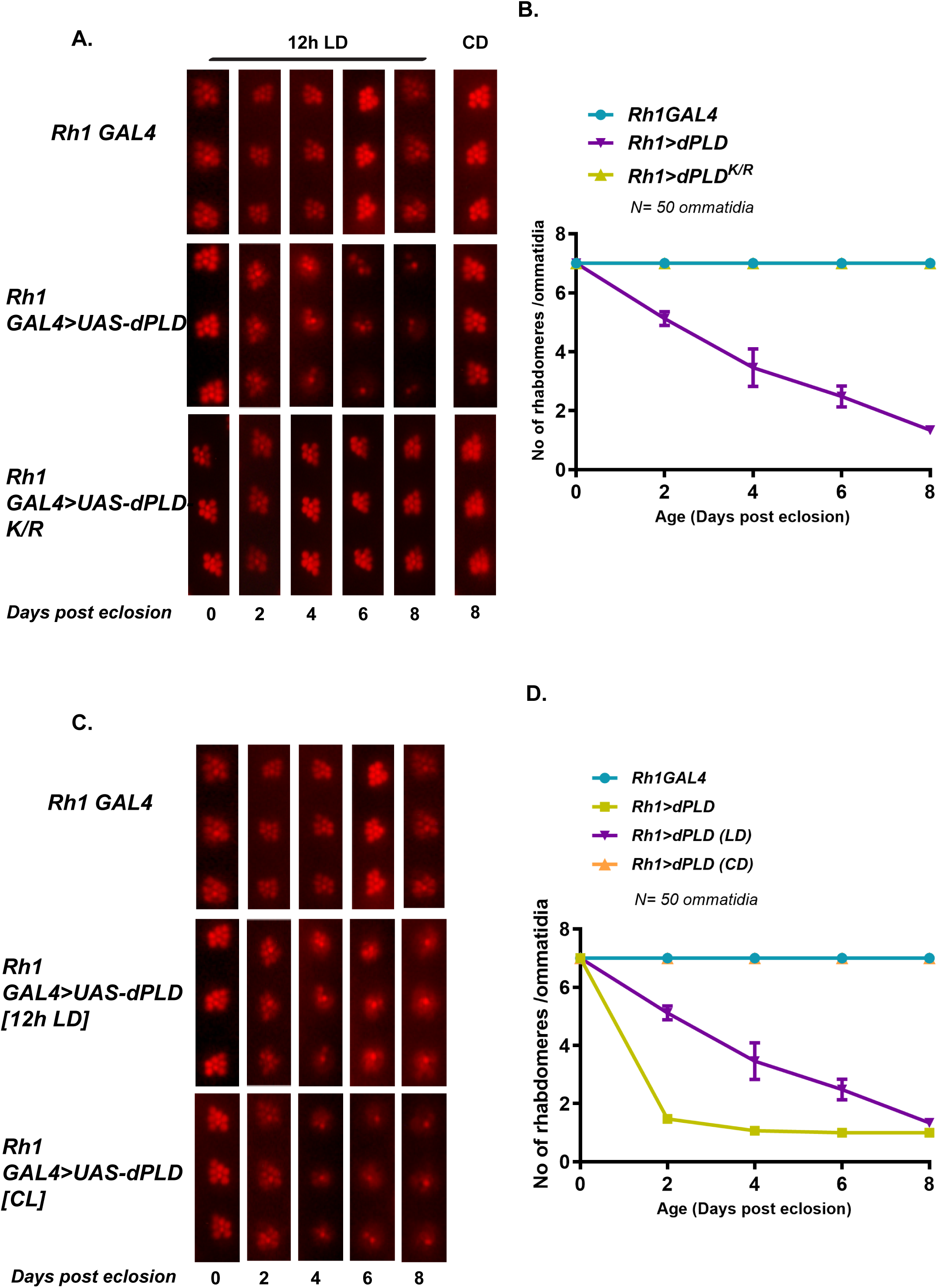
(A) Representative optical neutralization (ON) images showing rhabdomere structure from *control*, *Rh1>dPLD* and *Rh1>dPLD^K/R^*. The age and rearing conditions are mentioned at the bottom of the panels. (B) Quantification of rate of PR degeneration of *control*, *Rh1>dPLD* and *Rh1>dPLD^K/R^* reared in 12h LD cycle. The X-axis represents age of the flies and the Y-axis represents the number of intact rhabdomere visualized in each ommatidium. n= 50 ommatidia taken from at least five separate flies. (C) Representative optical neutralization (ON) images showing rhabdomere structure from *control* and *Rh1>dPLD*. The rearing conditions are mentioned on the left. The age is mentioned at the bottom of the panels. (D) Quantification of rate of PR degeneration of *control*, *Rh1>dPLD* reared in different light conditions. The X-axis represents age of the flies and the Y-axis represents the number of intact rhabdomere visualized in each ommatidium. n= 50 ommatidia taken from at least five separate flies.

### PX and PH domains regulate dPLD activity and localisation

Since the activity of many enzymes is often regulated by additional domains in the protein, we studied the role of the non-catalytic domains of dPLD in regulating its activity. As a readout of activity, we monitored the severity of retinal degeneration upon overexpressing dPLD. In addition to its catalytic HKD domains, dPLD also contains an N-terminal Phox homology (PX) and Pleckstrin homology (PH) domains that are well conserved with its mammalian orthologs (Raghu et al., 2009b).

To test the role of PX and PH domains in regulation of dPLD activity, we generated a construct where both PX and PH domains were deleted (*dPLD*^Δ*PXPH*^) [**Fig. 2A**]. In contrast to overexpression of dPLD, overexpression of dPLD^ΔPXPH^ did not lead to light dependent retinal degeneration suggesting that dPLD^ΔPXPH^ is not activated in response to illumination [**Fig. 2B, C**]. Further we reconstituted dPLD^ΔPXPH^ in *dPLD^3.1^*, a loss-of-function mutant of dPLD (Thakur et al., 2016) and tested if it could rescue the retinal degeneration phenotype of *dPLD^3.1^*. We found that dPLD^ΔPXPH^ was unable to rescue the retinal degeneration in *dPLD^3.1^*. [**Fig 2D**]. Under the same conditions wild type dPLD was able to rescued the phenotype of *dPLD^3.1^*. Immunofluorescence analysis revealed that the dPLD^ΔPXPH^ protein was distributed throughout the cell body in contrast to the wild type dPLD protein that is strictly localised to the base of the rhabdomere [**Fig. 2E**].

**Figure 2:**
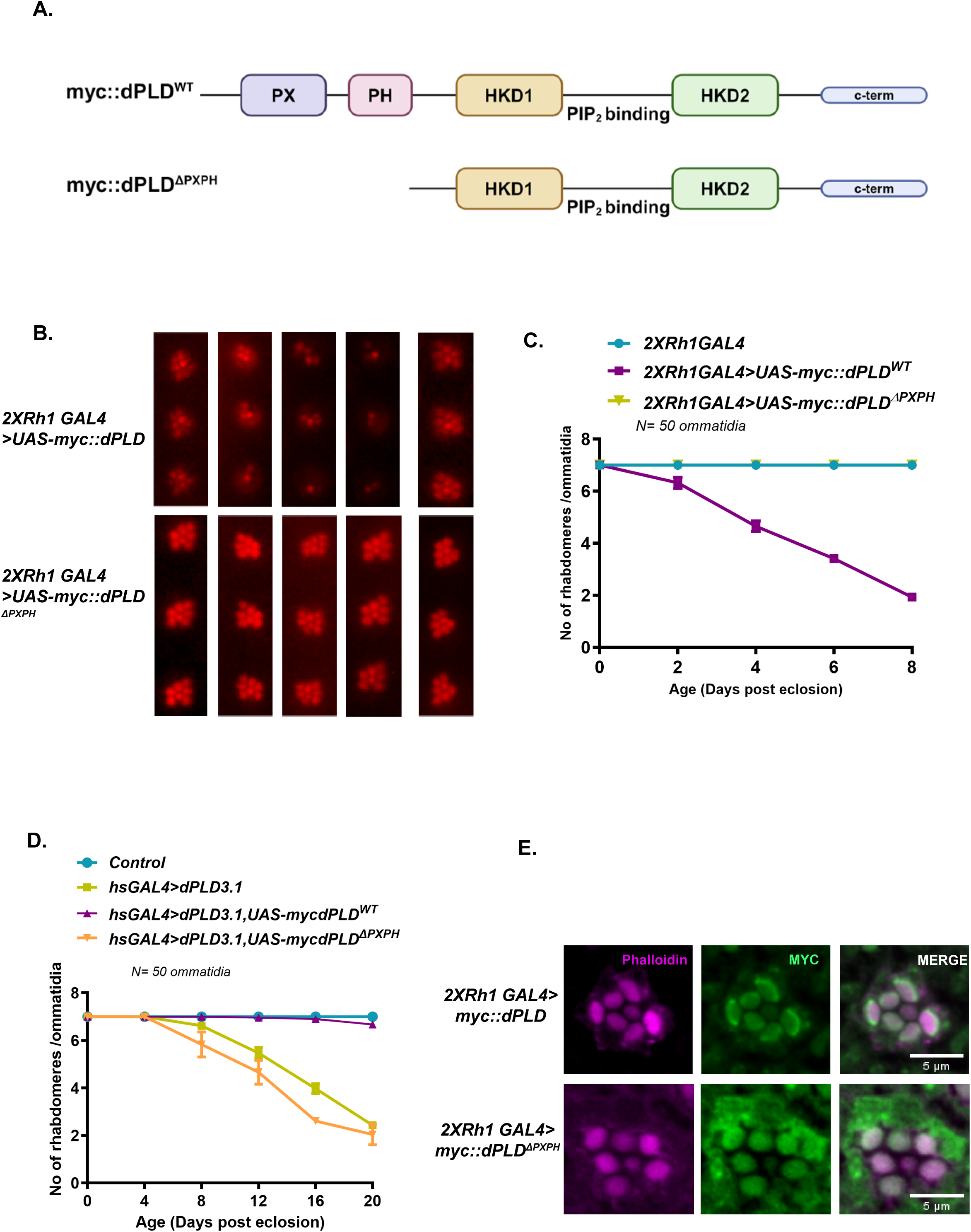
(A) Domain structure of myc::dPLD^WT^ and myc::dPLD^ΔPXPH^. (B) Representative optical neutralization (ON) images showing rhabdomere structure from 2X*Rh1>myc::dPLD* and *2XRh1> myc::dPLD^ΔPXPH^*. The age and rearing conditions are mentioned at the bottom of the panels. (C) Quantification of rate of PR degeneration of *control*, 2X*Rh1>myc::dPLD* and *2XRh1> myc::dPLD^ΔPXPH^* reared in 12h LD cycle. The X-axis represents age of the flies and the Y-axis represents the number of intact rhabdomere visualized in each ommatidium. n= 50 ommatidia taken from at least five separate flies. (D) Quantification of rate of PR degeneration of *control*, *hsGAL4,dPLD^3.1^, hsGAL4,dPLD^3.1^>myc::dPLD* and *hsGAL4,dPLD^3.1^> myc::dPLD*^Δ*PXPH*^ reared in CL. The X-axis represents age of the flies and the Y-axis represents the number of intact rhabdomere visualized in each ommatidium. n= 50 ommatidia taken from at least five separate flies. (E) Transverse section (TS) of retinae from 2X*Rh1>myc::dPLD* and *2XRh1> myc::dPLD*^Δ*PXPH*^stained with MYC antibody. Flies were dissected after 0-6 hrs (day 0) post eclosion.

### The PH domain is required for the correct localization of dPLD

To test the contribution of the PH domain to the localization and function of dPLD, we deleted this domain (*dPLD*^Δ*PH*^) [**Fig. 3A**]. Overexpression of *dPLD*^Δ*PH*^ showed no light dependent retinal degeneration [**Fig. 3B, C**] implying loss of dPLD activity. Likewise, reconstitution of dPLD^ΔPH^ was not able to rescue the retinal degeneration phenotype of *dPLD^3.1^* [**Fig 3D**]. Further, we found that dPLD^ΔPH^ was mislocalised like *dPLD*^Δ*PXPH*^ away from the base of the rhabdomere [**Fig. 3E**]. Thus, the PH domain is necessary for correct localisation of dPLD and activity in photoreceptors.

**Figure 3:**
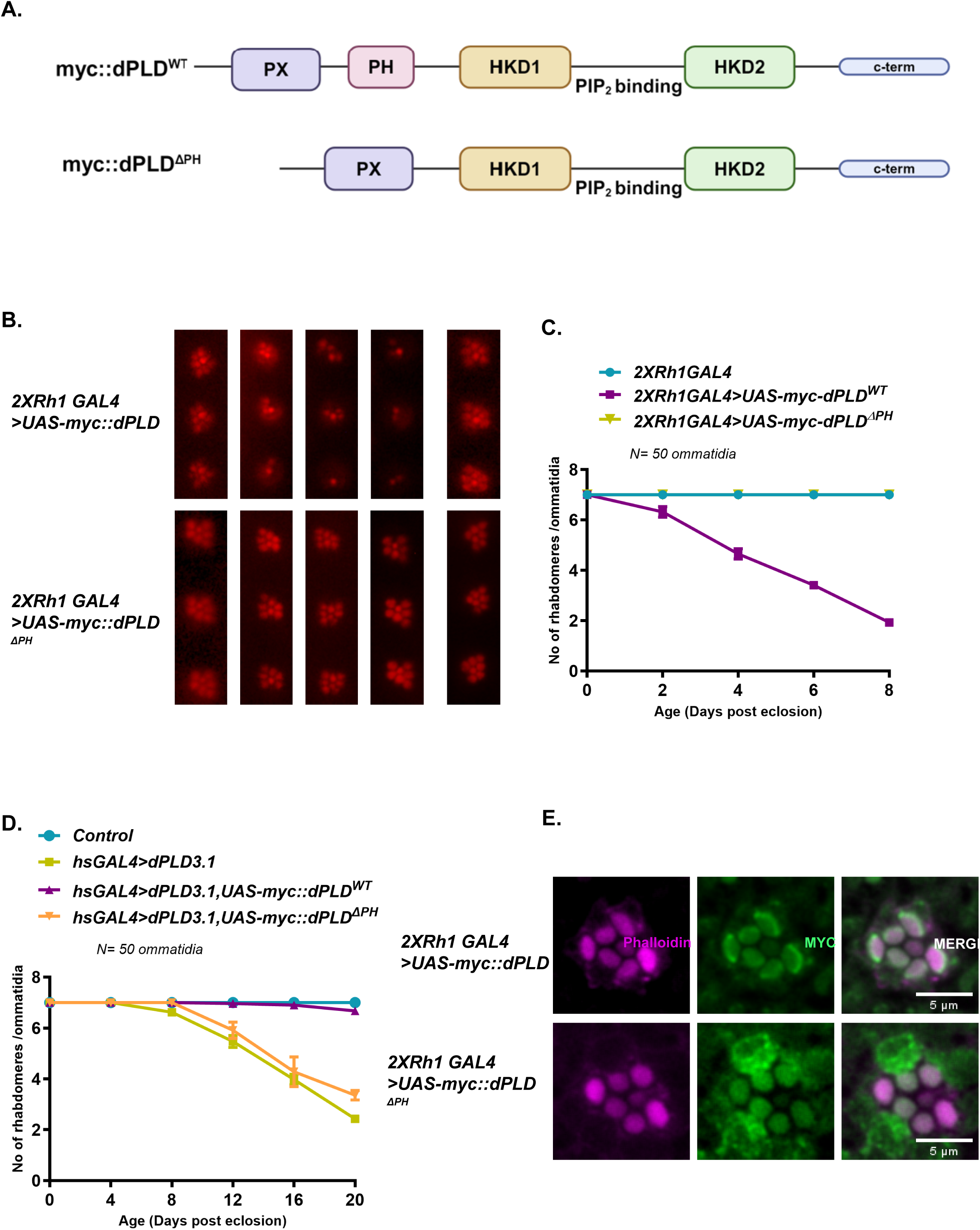
(A) Domain structure of myc::dPLD^WT^ and myc::dPLD^ΔPH^. (B) Representative optical neutralization (ON) images showing rhabdomere structure from 2X*Rh1>myc::dPLD* and *2XRh1> myc::dPLD*^Δ*PH*^. The age and rearing conditions are mentioned at the bottom of the panels. (C) Quantification of rate of PR degeneration of *control*, 2X*Rh1>myc::dPLD* and *2XRh1> myc::dPLD*^Δ*PH*^ reared in 12h LD cycle. The X-axis represents age of the flies and the Y-axis represents the number of intact rhabdomere visualized in each ommatidium. n= 50 ommatidia taken from at least five separate flies. (D) Quantification of rate of PR degeneration of *control*, *hsGAL4,dPLD^3.1^, hsGAL4,dPLD^3.1^>myc::dPLD* and *hsGAL4,dPLD^3.1^> myc::dPLD*^Δ*PH*^ reared in CL. The X-axis represents age of the flies and the Y-axis represents the number of intact rhabdomere visualized in each ommatidium. n= 50 ommatidia taken from at least five separate flies. (E) Transverse section (TS) of retinae from 2X*Rh1>myc::dPLD* and *2XRh1> myc::dPLD*^Δ*PH*^stained with MYC antibody. Flies were dissected after 0-6 hrs (day 0) post eclosion.

### The PX domain negatively regulates dPLD activity

The PX domain is known to regulate the function of many proteins through its ability to bind to phosphatidylinositol 3 phosphate (PI3P) (Xu et al., 2001). We tested the ability of dPLD to bind with PI3P using a lipid overlay assay and found that a concentration dependent increase in PI3P binding to dPLD *in vitro* [**Fig 4B, C**]. When compared to the wild type protein, dPLD^ΔPX^ **[Fig. 4A]** showed reduced binding to PI3P under the same experimental conditions [**Fig. 4B, C**], although PI3P binding was not abolished suggesting that although the PX domain of dPLD binds PI3P, there is another PI3P binding site in dPLD.

**Figure 4:**
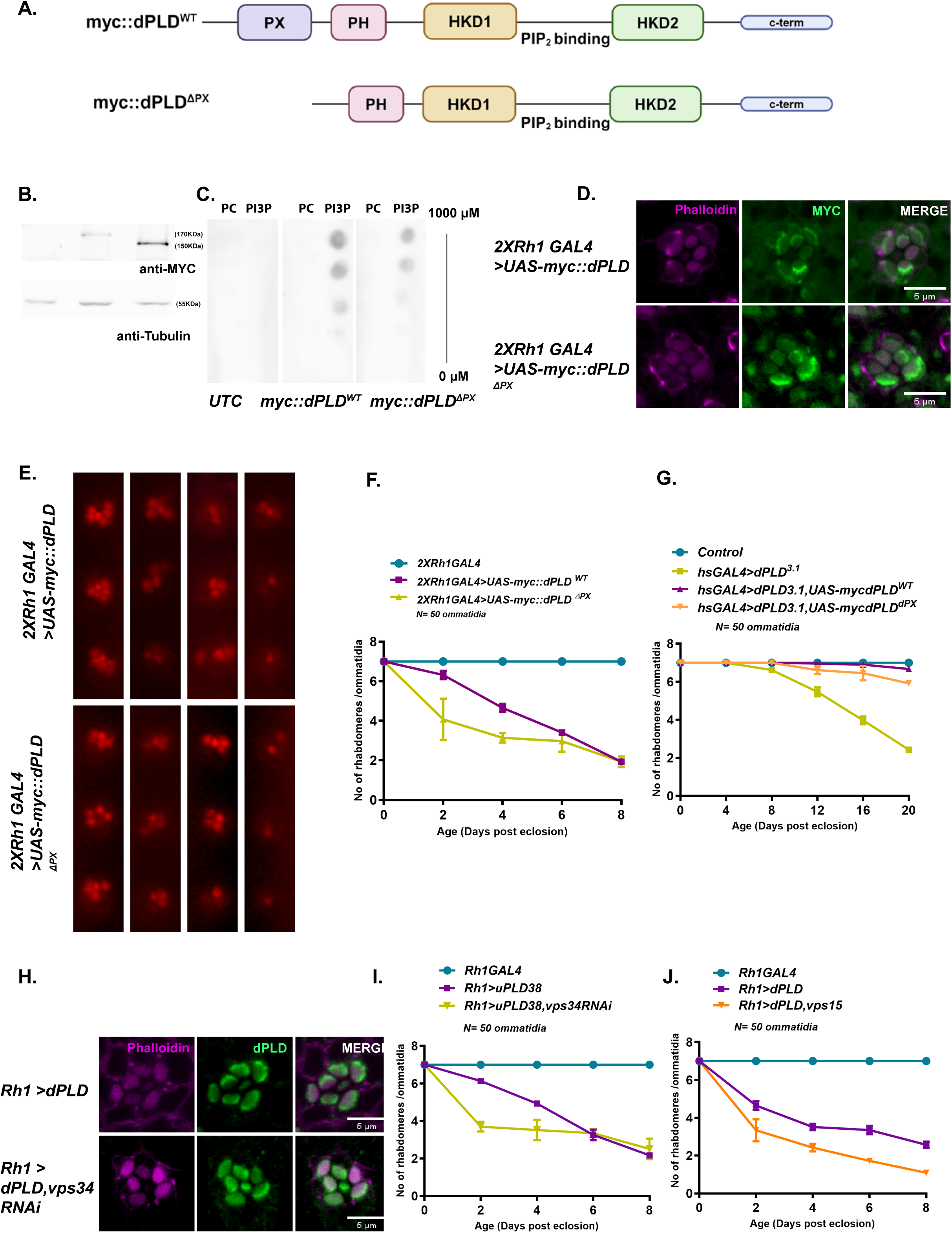
(A) Domain structure of myc::dPLD^WT^ and myc::dPLD^ΔPX^. (B)Western blot for expression of untransfected control, myc::dPLD^WT^ and myc::dPLD^ΔPX^ S2 cell lysate probed using myc antibody. Tubulin was used as loading control. (C) Lipid overlay assay using control and PI3P for untransfected and myc::dPLD^WT^ and myc::dPLD^ΔPX^ S2 cell lysates. The lipids are spotted in increasing concentrations from bottom to top. The strips were probed with MYC antibody. (D) Transverse section (TS) of retinae from 2X*Rh1>myc::dPLD* and *2XRh1> myc::dPLD*^Δ*PX*^ stained with MYC antibody. Flies were dissected after 0-6 hrs (day 0) post eclosion. (E) Representative optical neutralization (ON) images showing rhabdomere structure from 2X*Rh1>myc::dPLD* and *2XRh1> myc::dPLD*^Δ*PX*^. The age and rearing conditions are mentioned at the bottom of the panels. (F) Quantification of rate of PR degeneration of *control*, 2X*Rh1>myc::dPLD* and *2XRh1> myc::dPLD*^Δ*PX*^ reared in 12h LD cycle. The X-axis represents age of the flies and the Y-axis represents the number of intact rhabdomere visualized in each ommatidium. n= 50 ommatidia taken from at least five separate flies. (G) Quantification of rate of PR degeneration of *control*, *hsGAL4,dPLD^3.1^, hsGAL4,dPLD^3.1^>myc::dPLD* and *hsGAL4,dPLD^3.1^> myc::dPLD*^Δ*PX*^ reared in CL. The X-axis represents age of the flies and the Y-axis represents the number of intact rhabdomere visualized in each ommatidium. n= 50 ommatidia taken from at least five separate flies. (H) Transverse section (TS) of retinae from *Rh1>dPLD and Rh1>dPLD,vps34RNAi* stained with dPLD antibody. Flies were dissected after 0-6 hrs (day 0) post eclosion. (I) Quantification of rate of PR degeneration of *control*, *Rh1>dPLD, Rh1>vps34RNAi, Rh1>dPLD,vps34RNAi* reared in 12h LD cycle. The X-axis represents age of the flies and the Y-axis represents the number of intact rhabdomere visualized in each ommatidium. n= 50 ommatidia taken from at least five separate flies. (J) Quantification of rate of PR degeneration of *control*, *Rh1>dPLD, Rh1>vps15RNAi, Rh1>dPLD,vps15RNAi* reared in 12h LD cycle. The X-axis represents age of the flies and the Y-axis represents the number of intact rhabdomere visualized in each ommatidium. n= 50 ommatidia taken from at least five separate flies.

One possible role of the PX-domain PI3P interaction is the localization of proteins containing this domain to PI3P containing compartments. To test this, we determined the localization of dPLD and dPLD^ΔPX^; this revealed no change in the localization of dPLD^ΔPX^ compared to dPLD [**Fig. 4D**] suggesting PX domain is dispensable for the normal localisation of dPLD.

An alternative possibility is that the PX domain could regulate the activity of dPLD. To test this, we studied the effect of overexpressing *dPLD*^Δ*PX*^ in comparison to dPLD. This study revealed that overexpression of *dPLD*^Δ*PX*^ resulted in light dependent retinal degeneration which was faster than that seen on dPLD overexpression [**Fig. 4E, F**]. Lastly, we reconstituted *dPLD^3.1^* with *dPLD*^Δ*PX*^ and found that it could rescue the retinal degeneration phenotype of the null mutant **[Fig. 4G]**. Therefore, the PX domain is not required for dPLD activity but may act as a negative regulator.

### PI3P levels regulate dPLD function in photoreceptors

To understand the role of PI3P in regulation of dPLD activity, we tested the effect of depleting this lipid on dPLD activity. As in other eukaryotes, in *Drosophila*, PI3P is primarily synthesized by the class III PI3K vps34 (Linassier et al., 1997) that acts in a complex with vps15 and in the early endosomal compartment the UVRAG complex regulates its activity (Itakura et al., 2008; Lee et al., 2011). Depletion of vps34 did not change the localization of dPLD [**Fig. 4H**]. However, depletion of vps34 by two independent RNAi lines while overexpressing dPLD (*Rh1>dPLD, vps34RNAi*) resulted in enhanced degeneration kinetics compared to dPLD overexpression (*Rh1>dPLD*) alone [**Fig. 4I**]. Independently, we knocked down another component of the vps34 complex (vps15) that should affect the function of the complex, resulting in reduced PI3P levels and phenocopy knockdown of vps34. Independent depletion of vps15 in dPLD overexpression background also resulted in enhanced degeneration kinetics phenocopying the effect of vps34 depletion [**Fig. 4J**]. This implies vps34 and in turn PI3P is a negative regulator of dPLD activity

### dPLD shows vesicular localisation and is regulated at the early endosomal compartments

Our data showing a role for PI3P in regulating dPLD activity suggests that PLD might be localized to compartments enriched in this lipid. It has been previously reported in photoreceptors, dPLD is localised at the base of the rhabdomere (LaLonde et al., 2005; Raghu et al., 2009a). However, PI3P is not enriched at the base of the rhabdomere [**Fig. 5A**]. Therefore, we imaged photoreceptors expressing dPLD to test if a pool of vesicular dPLD exists. We noted that along with the previously reported (Lalonde, Raghu JCB) enrichment at the base of the rhabdomere, there is also a vesicular pool of dPLD [**Fig. 5B**].

**Figure 5:**
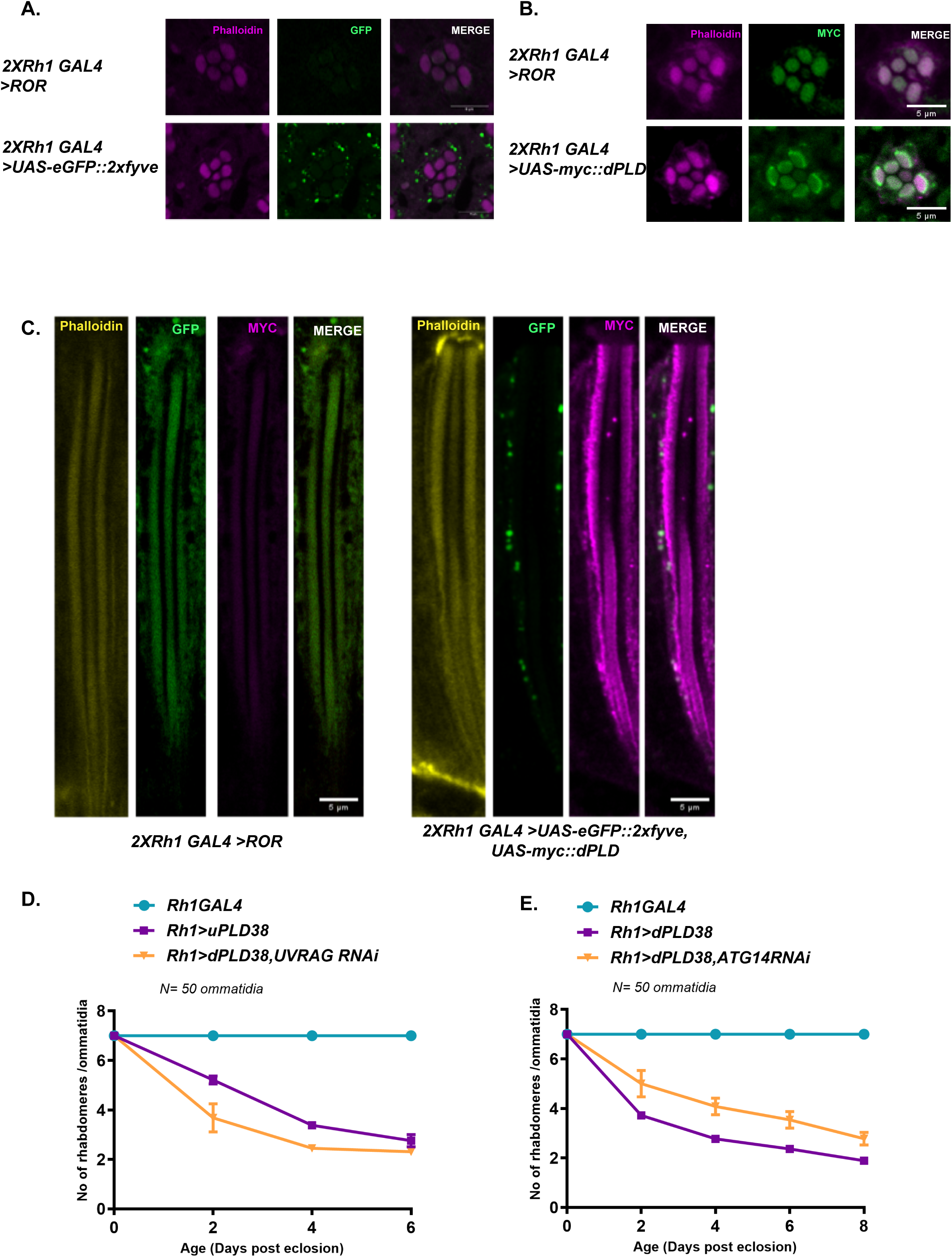
(A) Transverse section (TS) of retinae from *control* and *Rh1>2xFYVE::eGFP* stained with GFP antibody. Flies were dissected after 0-6 hrs (day 0) post eclosion. (B) Super-resolution confocal imaging for transverse section (TS) of retinae from *WT* and 2X*Rh1>myc::dPLD* stained with MYC antibody. Flies were dissected after 0-6 hrs (day 0) post eclosion. (C) Longitudinal section (LS) of retinae from *2XRh1> ROR and 2XRh1>2xFYVE::eGFP,myc::dPLD* stained with MYC and GFP antibody. Flies were dissected after 0-6 hrs (day 0) post eclosion. (D) Quantification of rate of PR degeneration of *control*, *Rh1>dPLD, Rh1>UVRAGRNAi, Rh1>dPLD,UVRAGRNAi* reared in 12h LD cycle. The X-axis represents age of the flies and the Y-axis represents the number of intact rhabdomere visualized in each ommatidium. n= 50 ommatidia taken from at least five separate flies. (E) Quantification of rate of PR degeneration of *control*, *Rh1>dPLD, Rh1>ATG14RNAi, Rh1>dPLD,ATG14RNAi* reared 14h LD cycle. The X-axis represents age of the flies and the Y-axis represents the number of intact rhabdomere visualized in each ommatidium. n= 50 ommatidia taken from at least five separate flies.

We studied the identity of the vesicular pool of myc::dPLD in photoreceptors. A fraction of myc::dPLD showed colocalization with PI3P compartments marked by the 2XFYVE probe [**Fig. 5C**] where as a proportion of dPLD did not localize with PI3P enriched compartments. To test which pool of PI3P was relevant for regulation of dPLD activity, we independently depleted UVRAG and ATG14 to individually deplete the early endosomal and autophagic pools of PI3P respectively. Knockdown of UVRAG (*Rh1>dPLD, UVRAG RNAi*) resulted in an enhancement of retinal degeneration in dPLD overexpressing photoreceptors, like the effect of vps34 depletion [**Fig 5D**]. By contrast, depletion of ATG14 in dPLD overexpression background (*Rh1>dPLD, ATG14 RNAi*) resulted in a slowing of retinal degeneration [**Fig 5E**].

## Discussion

Many studies over the years have documented that PLD is an enzyme activated in the context of ligand binding to G-protein coupled receptors and, in many cases, this occurs alongside phospholipase C activation. However, the mechanism and location at which PLD is activated has not been well understood. In mammalian systems there are typically two PLD activities, PLD1 localised to a vesicular compartment within cells and PLD2 that is found at the plasma membrane, complicating the analysis of this problem. By contrast, in the *Drosophila* genome, there is only a single PLD (LaLonde et al., 2005; Raghu et al., 2009a) and previous work has shown that its localization and function resembles that of mammalian PLD1 (Panda et al., 2018; Raghu et al., 2009b) In adult *Drosophila* photoreceptors, dPLD is reported to be localised very specifically to the base of the microvillar plasma membrane, most likely at the sub-microvillar cisternae (SMC) that are a specialised sub-compartment of the smooth endoplasmic reticulum (Yadav et al., 2016) and in these cells, dPLD is activated in the context of phototransduction (Thakur et al., 2016), a process where rhodopsin, a G-protein coupled receptor activates phospholipase C (Raghu et al., 2012). Using this model system, we found that overexpression of dPLD results in a light dependent retinal degeneration that is also dependent on the lipase activity of the enzyme. In this setting, we find that accurate localization of dPLD is important for its activity. Deletion of the PH domain (*dPLD*^Δ*PH*^) mislocalizes the enzyme away from the region of the SMC along with loss of its function as reported by the retinal degeneration assay. This loss of function may be a consequence either of loss of enzymatic activity of *dPLD*^Δ*PH*^, for.e.g due to loss of co-localization with an activator, or due to dPLD activity now being at an alternate subcellular location to where it is required. Although we find that the PH domain is essential for localization to the SMC, the ligand with which the PH domain interacts remains to be identified. Our finding that the PH domain of dPLD is required for its localization is consistent with studies on mammalian PLD1 where the PH domain has been shown to be important for localization and fatty acylation of residues in the PH domain has also been shown to be a key determinant (Sugars et al., 2002, 1999).

In this study, we found that reconstitution of a null mutant of dPLD in *Drosophila* with *dPLD*^Δ*PH*^ could not rescue its phenotypes, implying that the localization of the wild type enzyme to the region of the SMC is important for supporting photoreceptor function *in vivo.* dPLD activity produces PA and our findings must imply that PA production by dPLD during phototransduction is important in the region of the SMC. Interestingly, diacylglycerol kinase (DGK) encoded by *rdgA*, has also been localized to this part of the photoreceptor (Masai et al., 1997) although *rdgA* loss of function mutants (Harris and Stark, 1977; Raghu et al., 2000) show a distinctive phenotype to dPLD mutants (Thakur et al., 2016). This observation suggests that the PA produced by dPLD and DGK support distinct processes in *Drosophila* photoreceptor structure and function. One possible biochemical basis for this model is the proposal that DGK and PLD produce PA species with distinctive acyl chains (Hodgkin et al., 1998). Therefore, although produced at the same sub-cellular location, PA produced by each of the two enzymes may have distinct cellular targets and functions.

In contrast to the PH domain, we found that the PX domain, a well know PI3P binding module is not required for accurate localization. Secondly, we could not demonstrate an enrichment of PI3P at the base of the microvilli, the location where dPLD is localized. Lastly, depletion of PI3P levels through down regulation of the vps34 complex, the key Class III PI3K enzyme regulating PI3P levels did not affect the localization of dPLD. Thus, overall, our findings imply that dPLD is localized to the SMC region via a PI3P independent mechanism that requires the presence of its PH domain. Although we found no role for PI3P binding in the localization of dPLD, we noted that the enzyme binds PI3P through both the PX domain as well as a region of the protein outside this domain. What is the function of this PI3P binding? We found that reconstitution of a null mutant of dPLD with *dPLD*^Δ*PX*^ *was* able to largely rescue the retinal degeneration phenotype of the mutant, something that depends on the catalytic activity of dPLD (this study). This finding suggests that the PX domain is dispensable for dPLD activity. However, we also found that overexpression of dPLD^ΔPX^ resulted in a retinal degeneration phenotype stronger than that resulting from dPLD overexpression. This finding may suggest that the PX domain is a negative regulator of dPLD activity. Consistent with this model, we also found that depletion of either vps34 and vps15, both subunits of the Class III PI3K complex also resulted on an enhancement of retinal degeneration resulting from dPLD overexpression. Taken together, these observations are consistent with a role for PI3P dependent negative regulation of dPLD activity. However, it has also been reported that the PX domain of mammalian PLD1 binds PI(3,4,5)P_3_,PA and PS (Lee et al., 2005; Stahelin et al., 2004) the role of these lipids in interacting with the PX domain of dPLD remains to be investigated.

Overall, this study assigns *in vivo* functions to two of the non-catalytic domains of dPLD; while the PH domain is required for the correct localization of dPLD, the PX domain appears to negatively regulate activity *in vivo*. The molecular mechanisms by which this regulation occurs remains to be elucidated.

## Acknowledgments

This work was supported by the Department of Atomic Energy, Government of India, under Project Identification No. RTI 4006, We thank the NCBS Imaging Facility, DNA sequencing and *Drosophila* facility for support.

## Materials and Methods

### Fly stocks and rearing conditions

All flies (*Drosophila melanogaster*) were reared on enriched media (composition:80g Cornflour, 20g D-Glucose, 40g Sugar, 8g Agar, 15g Yeast, 4ml Propionic acid, 0.7g Tego, 0.6ml Orthophosphoric acid for 1 litre of media) in 25°C incubators with no internal illumination. For experiments involving exposure to illumination, the flies were reared in incubators with illumination conditions of wavelength: 400-700nm; intensity: ∼1500 lux with either constant light or 12h light-dark cycle. For constant dark rearing, fly vails were kept in a closed dark box in 25°C incubator with no internal illumination for the specified time.

Wild type (WT) used were Oregon-R. The GAL4-UAS system (Brand and Perrimon, 1993) was used to express transgenic constructs. The following transgenic lines were obtained from the Bloomington Stock Center: UAS-vps34 RNAi (BL33384), UAS-vps34 RNAi (BL64011), UAS-UVRAG RNAi (BL 34368), UAS-ATG14 RNAi (BL 40858).

### Generation of transgenic lines

Site-specific integration (Fly Facility, NCBS, Bangalore) at an attP2 site on the third chromosome was used to generate transgenic flies.

### Optical Neutralization and Scoring Retinal Degeneration

Flies were immobilized by cooling on ice and decapitated with a sharp blade. The fly head was fixed on a glass slide using a drop of colourless nail varnish. The refractive index of the cornea was neutralized using a drop of immersion oil and imaging was done using 40X oil objective of Olympus BX43 microscope. The digital image acquisition and documentation were done by using CellSens software.

To calculate the photoreceptor degeneration index, five flies per time point were scored. A total of 50 ommatidia were used per time point to calculate the photoreceptor degeneration index. A score of 1 was given to each rhabdomere that looked like wild type. Mutants with degenerated photoreceptors will have a score between 1 and 7. Degeneration index were represented as mean ± SEM.

### Immunohistochemistry

Retinae from adult flies were scooped out in ice cold PBS under low red light and then fixed in 4% paraformaldehyde with 1mg/ml saponin for 15 min at room temperature. Fixed eyes were washed thrice with PBTX (PBS with 0.3% TritonX-100). Retinae were blocked in solution containing 10% fetal bovine serum (FBS) in PBTX for 2 hours at room temperature, after which they were incubated with primary antibodies diluted in blocking solution. Primary antibodies used were against Rh1, 1:50(4C5-C, DSHB), myc, 1:200(2272S CST), GFP, 1:2000(ab13970 Abcam Cambridge, UK), EEA1, 1:500 (Mario Zerial, MPI-CBG, Dresden, Germany) and Drosophila ATG8a,1:500 (Rachel Kraut, Biological Sciences, Nanyang Technological University, Singapore) for 24 hours at 4°C on a shaker. Appropriate secondary antibodies conjugated with a fluorophore were used at 1:300 dilutions [Alexa Fluor 633/568/488 IgG, (Molecular Probes)] and incubated for 8 hours at 4°C on a shaker. Wherever required, phalloidin conjugated to Alexa-flours was incubated along with secondary antibodies to stain for F-actin. Post this, the tissues were washed with PBTX followed by PBS and mounted in 70%glycerol in PBS. These whole-mounted preparations were imaged under a 60X 1.4NA oil objective, in an Olympus FV3000 laser scanning confocal microscope.

### Lipid overlay assay

#### Cell culture

S2R+ cells stably expressing Actin>GAL4 were grown in Schneider’s Drosophila Insect Medium (SDM + L-Glutamine, Invitrogen) supplemented with 10% heat inactivated fetal bovine serum (FBS, Gibco-BLR), 1ug/ml Puromycin and 1:100 of Penicillin-Streptomycin solution (Gibco)- (SCM-Schneider’s Complete Medium, at 25°C.

#### Transfection and sample preparation

S2R+ cells were individually transfected with pUAST-myc::dPLD-attB, pUAST-myc::dPLDΔPX -attB using Effectene transfection kit (Qiagen) following manufacturer’s protocol for 36 h. Following which cells were pelleted at 1000 rpm and washed twice with PBS. The final pellet was lysed in FAT BLOT buffer (50mMTris-Cl pH 7.5, 150MNaCl) containing PIC.

#### Lipid strips

In parallel, lipid strips made using nitrocellulose membranes [Hybond-C Extra; (GE Healthcare, Buckinghamshire, UK)] spotted with increasing picomoles (0-1000uM) of DOPC (Avanti Polar Lipids, 850375C) and PI3P (Eschelon Biosciences, P-3016). The spotted membrane was dried for about 30 mins at room temperature and then blocked using 5% BSA (HiMedia) in FAT BLOT buffer for 2 h at RT. Following this, the strips were incubated overnight at 4°C with the respective cell lysates in blocking solution (FAT-BLOT buffer+5% Blotto + 0.06% Tween20) such that the amount of overexpressed protein is equal across different samples (Quantified using Bradford assay and western blotting). Next the membranes were washed extensively three times with FAT BLOT buffer containing 0.1% Tween20 (Sigma-Aldrich) [used for all subsequent wahes] and then incubated with anti myc (1:1000, 2272S CST) diluted in FAT BLOT buffer containing 5%Blotto and 0.1% Tween20 at RT for 3 h. After 3 washes the membranes were then probed with the appropriate HRP-conjugated secondary antibody (Jackson Immunochemicals; 1:10,000) in FAT BLOT buffer containing 5%Blotto and 0.1% Tween20. Following 3 washes, the binding was detected using ECL (GE Healthcare) in a LAS4000 instrument.

### Western blot

For retinal extract preparation, the flies were snap frozen in liquid nitrogen, followed by Acetone fixation and dehydration at −80°C for 2 days. Then the acetone was drained off and was allowed to completely vaporize off form the tissues in next 2days at RT. Post this the retinae were dissected out from the heads using a clean scalpel blade and were homogenised in 2X SDS PAGE sample buffer (with 4% SDS) followed by incubation at 95°C for 5mins.

The samples were subjected to separation using SDS-PAGE and the proteins were transferred to a nitrocellulose membrane [Hybond-C Extra; (GE Healthcare)] using wet transfer apparatus (Bio-Rad). The membrane was blocked using 5% Blotto (sc-2325, Santa Cruz Biotechnology) in phosphate buffer saline (PBS) with 0.1% Tween 20 (Sigma-Aldrich) (PBST) for 2 h at room temperature (RT). Primary antibody incubation was done overnight at 4°C using appropriate antibody dilutions: anti-myc (1:1000, 2272S CST), anti-α-tubulin (1:4000, E7c DSHB). Following this, the membrane was washed with PBST and incubated with 1:10,000 dilutions of appropriate secondary antibody (Jackson ImmunoResearch Laboratories) conjugated to horseradish peroxidase in blocking solution at RT for 2 h. After three PBST washes, the blots were developed with ECL (GE Healthcare) and imaged using LAS 4000 instrument (GE Healthcare).

### Molecular biology

The Pld cDNA used in this study originated from the clone pOT2a-GH07346 generated by the Berkeley Drosophila Genome Project. This clone has been fully sequenced by the Berkeley Drosophila Genome Project, and its sequence is available in GenBank (accession no. AF145640). It was amplified with myc tag at its N-terminal and cloned into a PUAST-attB vector using Not1 and Xba1 restriction enzymes (NEB). This was used as the parent vector for generation of all further domain deletion constructs. For dPLDΔPXPH, the parent vector was back amplified deleting the region corresponding to amino acids spanning 159-492 which comprises PX and PH domains. The amplicon was phosphorylated at the ends using PNK reaction and the ends were ligated using T4 DNA ligase (NEB). Simultaneously individual domain deletion constructs, dPLD^ΔPX^ (159-361 amino acids) and dPLD^ΔPH^ (371-492 amino acids) were also generated using the same method.

For cloning of PUAST-eGFP-2X- FYVE-attB, 2X-FYVE was cloned into pUAST-attB vector using EcoR1 and Xba1 restriction enzymes. eGFP was added to the N terminal region along with a gly-ser linker using GIBSON assembly.

